# Timing-dependent effects of elevated maternal glucocorticoids on offspring brain gene expression in a wild small mammal

**DOI:** 10.1101/2024.04.19.590319

**Authors:** Sarah E. Westrick, Eva K. Fischer, Freya van Kesteren, Stan Boutin, Jeffrey E. Lane, Andrew G. McAdam, Ben Dantzer

**Author notes:** **Corresponding author:** Sarah E Westrick.

## Abstract

An increase in maternal stress during offspring development can have cascading, life-long impacts on offspring behavior and physiology, which can vary depending on the timing of exposure to the stressor. By responding to stressors through increasing production of glucocorticoids (GCs), the hypothalamic-pituitary-adrenal (HPA) axis is a key mediator of maternal effects – both on the side of the mother and the offspring. At a molecular level, maternal effects are thought to be mediated through modifying transcription of genes, particularly in the brain. To better understand the evolutionary implications of maternal effects, more studies are needed on mechanisms of maternal effects in wild populations. To test how the timing of maternal stress impacts gene expression in the brains of offspring, we treated free-ranging North American red squirrels (*Tamiasciurus hudsonicus*) with GCs during late pregnancy or early lactation and collected brains from offspring around weaning. We used RNA-sequencing to measure gene expression in the hypothalamus and hippocampus. We found small differences in gene expression between GC-treated and control individuals suggesting long-term effects of the GC treatment on neural gene transcription. The general patterns of gene regulation across the transcriptome were consistent between the pregnancy and lactation-treated individuals. However, the number of significantly differentially expressed genes was higher in the lactation treatment group. These results support the idea that maternal stress affects neural gene expression in offspring, and these effects are dependent on timing. Our findings add valuable insight into the impact of maternal hormones on neural transcriptomics in a wild population.

## Introduction

The early life environment can have major downstream effects on offspring physiology and behavior, and ultimately fitness (Bateson et al., 2004; West-Eberhard, 2003). The parental environment and parental phenotypes, in particular, provide information that influences the development of many life-long phenotypes in offspring, thus shaping phenotypes across multiple generations (Badyaev & Uller, 2009). Importantly, the timing of when offspring receive cues can be an important factor in the resulting changes in phenotypes (Bell & Hellmann, 2019; McNamara et al., 2016). Developmental plasticity plays a critical role in evolution as it allows individuals to respond to information received in early life when many phenotypes can still be shifted adaptively along their developmental trajectories (West-Eberhard, 2003). Through multigenerational developmental plasticity, parental effects have the potential to impact multiple levels of biological organization including population and community dynamics (Badyaev & Uller, 2009; Duckworth et al., 2015; Rossiter, 1996; Snell-Rood et al., 2010).

Through behavioral and physiological interactions between mother and offspring, maternal effects in particular are considered one of the most important influences shaping the development of life-long offspring phenotypes (Bernardo, 1996; Mousseau & Fox, 1998; Wade, 1998). These maternal effects can have profound consequences for both mothers and their offspring (Maestripieri & Mateo, 2009). Maternal stress, in particular, has been found to have both adaptive and maladaptive effects depending on ecological context (Sheriff et al., 2017; Sheriff & Love, 2013). For example, maternal stress can be helpful if it prepares offspring for the environmental conditions they will face during development and as reproductive adults (i.e. predictive adaptive responses, Bateson et al., 2014), but also maladaptive when there is a mismatch between the maternal environment and offspring environment (Sheriff et al., 2018). While many biomedical studies are primarily focused on the pathological effects of perinatal stress (e.g. Weinstock, 2005), studies in evolutionary ecology often focus on the adaptive value of maternal stress (e.g. O. P. Love & Williams, 2008). In mammalian species, mothers play an especially vital role in shaping the early life environment. Mammalian mothers carry their offspring throughout early development and provide an extended period of care through lactation after birth (Maestripieri & Mateo, 2009; Reinhold, 2002; D. Roff, 1998).

From a proximate perspective, the maternal environment influences the developmental programming of many physiological processes in offspring including crucial neuroendocrine systems (Adkins-Regan et al., 2013; Champagne, 2013), such as the hypothalamic-pituitary-adrenal (HPA) axis (e.g. Meaney, 2001), hypothalamic-pituitary-gonadal (HPG) axis (e.g. Cameron, 2011), hypothalamic-pituitary-thyroid (HPT) axis (Hellstrom et al., 2012), oxytocinergic system (e.g. Champagne et al., 2001), serotonergic system (Courtiol et al., 2018), and dopaminergic system (Peña et al., 2019). These neuroendocrine systems act separately and in conjunction to modulate behavior such as the behavioral stress response (Barha et al., 2007; Weaver et al., 2004), social and sexual behavior (Cameron et al., 2008), learning and memory (Liu et al., 2000), and maternal care of the next generation (Champagne et al., 2001; Francis et al., 1999; Gonzalez et al., 2001). At a molecular level, changes in neuroendocrine systems due to the maternal environment are often underpinned by modulation of gene expression patterns (Champagne, 2008; Meaney, 2001; Murgatroyd et al., 2009; Parel & Peña, 2022; Peña et al., 2019; Weaver et al., 2002, 2004). For example, in lab mice, early life stress through maternal separation and limited nesting material can alter transcriptomic patterning such that offspring have an increased sensitivity to stressors as adults, as measured by behavioral responses to stressors (Peña et al., 2019). Overall, changes in brain gene expression shape neuroendocrine systems which can alter many downstream phenotypes.

The HPA axis is a particularly influential player in maternal effects, both on the side of the mother and also the offspring (Barbazanges et al., 1996; Glover et al., 2010; Kapoor et al., 2006; Maccari et al., 1995; Weinstock, 2008). In addition to its diverse roles in metabolism and the immune system, the HPA axis is responsible for modulating circulating glucocorticoids (GCs) in response to stressful events and environmental conditions (Charmandari et al., 2005; Sapolsky et al., 2000; Tsigos & Chrousos, 2002). GCs have many pleiotropic effects and are involved in many physiological processes (MacDougall-Shackleton et al., 2019). Like other steroid hormones, GCs and their receptors directly modulate transcription, thereby increasing or decreasing expression of target genes (Beato, 2000; Gray et al., 2017; Grbesa & Hakim, 2017) and also exert rapid nongenomic effects through second messenger cascades (Borski, 2000; Falkenstein et al., 2000; Jiang et al., 2014; Johnstone et al., 2019).

In mammals, GCs are passed from mother to offspring *in utero* across the placenta (Fowden & Forhead, 2015) and through milk (Grosvenor et al., 1993; Melo, 2015; Stead et al., 2022). Maternal GCs are generally higher in the mother than the fetus but the majority of the GCs in fetal circulation are of maternal origin (Hennessy et al., 1982; Huang et al., 2012). Maternal GCs during pregnancy and lactation provide valuable information to developing offspring about the conditions they will face (Fowden & Forhead, 2009, 2015). For example, higher concentrations of GCs in milk may physiologically “program” offspring to prioritize growth (Hinde et al., 2015). This programming is hypothesized to be the result of epigenetic modulation of the production of hormone receptors through changes in gene expression (Barbazanges et al., 1996; Champagne, 2013; Champagne & Meaney, 2006; Kapoor et al., 2006; Levine, 1994).

However, the impact of maternal GCs on development is often studied through experimentally elevating “early life stress” through techniques such as neonatal maternal deprivation/separation and exposing pregnant mothers to environmental stressors. This technique is valuable in probing the overall impact of maternal “stress” but can alter more variables than just maternal GCs, such as changes in catecholamines and arginine vasopressin (Charmandari et al., 2005). Direct manipulations of maternal GCs may be more appropriate for causally linking these “early life stress” effects to elevations in maternal GCs across development (e.g. Barbazanges et al., 1996).

Importantly, the effects of early life stress on offspring behavior may vary depending on the timing, both across (i.e., pre-vs postnatal) and within (i.e., early vs late lactation) developmental periods. Many laboratory studies have found that prenatal stress causes changes in HPA axis function by reducing HPA axis negative feedback, thus prolonging the release of glucocorticoids following exposure to a stressor (Caldji et al., 2000, 2011; Francis et al., 1999, 2002). However, results from studies on postnatal stress in rodents are more equivocal, which may be due to variation in timing of the postnatal stressor relative to the “stress hyporesponsive period”, an adrenal hyporesponsive period where the HPA axis shows a markedly reduced response (Schmidt, 2019). For example, in lab rats, early life stress by maternal separation impacts offspring behavior and immune functions differently depending on how much the timing of separation overlaps with this hyporesponsive period (Roque et al., 2014).

Despite strong implications for ultimate evolutionary consequences (Maestripieri & Mateo, 2009; Mousseau & Fox, 1998), the vast majority of mechanistic studies on mammalian maternal effects have been conducted in a controlled laboratory environment on traditional model organisms (typically mice and rats). The emerging field of ecological epigenetics is aimed at addressing this gap in knowledge (Bossdorf et al., 2008; Ledon-Rettig et al., 2013). Since phenotypes are shaped through the integration of multivariate cues from the environment (Dall et al., 2015), the increased complexity of natural systems may impact how maternal effects manifest. While controlled laboratory experiments can tell us whether or not a manipulation can have a measurable impact, these effects may not translate to natural populations where the environments, genetic backgrounds, and therefore phenotypes of individuals are often more variable. Additionally, with more information about the mechanisms underlying maternal effects, we can investigate how evolutionary trajectories are constrained or biased. There may be many different mechanistic routes to the same phenotype being measured, but which mechanisms are used and how “selectable” they are can constrain evolutionary trajectories (Hofmann et al., 2014; Rittschof & Robinson, 2014; D. A. Roff, 2007). Neuroendocrine systems, in particular, play an important pleiotropic role in correlational selection and may promote or constrain the effects of selection (McGlothlin & Ketterson, 2008). Thus, improving our understanding of the mechanisms underlying developmental and behavioral plasticity in natural populations will further inform our understanding of the evolutionary impact of maternal effects (Badyaev & Uller, 2009; Groothuis & Schwabl, 2007) and their role as animals face increased environmental stressors (reviewed in Meylan et al., 2012).

Small mammals, such as North American red squirrels (*Tamiasciurus hudsonicus*), provide excellent models for studying maternal effects due to comparably short generation times and ease of monitoring large numbers of individuals and their offspring (Dantzer et al., 2022). Red squirrels are also amenable to large-scale individual manipulations (Dantzer et al., 2013). The specific population of red squirrels in this study experiences annually fluctuating variation in food resources, competition, and predation risk (Dantzer, McAdam, et al., 2020). Among year variation in food availability due to masting of their primary food resource (seeds from white spruce [*Picea glauca*] trees) is closely mirrored by fluctuations in red squirrel population density (Dantzer et al., 2013; Dantzer, McAdam, et al., 2020). Red squirrels can anticipate years of high food availability and produce more offspring resulting in higher competition for territory acquisition (Boutin et al., 2006; Petrullo et al., 2023). If offspring can use cues from their mother to infer environmental conditions during development, they may be able to adaptively adjust physiological and behavioral phenotypes in preparation (Bateson et al., 2014; Dantzer, 2023). In fact, red squirrel pups born to mothers who experience perceived high conspecific density while pregnant adaptively grow faster than offspring from mothers without the cue of increased density (Dantzer et al., 2013). This adaptive response is likely driven by elevated maternal GCs during years of high conspecific density or perceived high conspecific density (Dantzer et al., 2013; Guindre-Parker et al., 2019). We have previously found that elevation in maternal GCs during pregnancy also results in faster growing pups (Dantzer et al., 2013; Dantzer, van Kesteren, et al., 2020) which may improve overwinter survival of pups (Hendrix et al., 2020; McAdam & Boutin, 2003), especially those encountering high densities after birth (Dantzer et al., 2013).

To further investigate the mechanisms of maternal effects in a wild population, we used a direct manipulation of maternal GCs at different timepoints in development. We experimentally elevated maternal GCs by individually supplementing pregnant or lactating mothers with exogenous GCs provisioned in a palatable food that the squirrels readily consume. From this manipulation, we discovered that the effects of elevated maternal GCs on offspring behavior and physiology varied depending on the timing of the manipulation. Offspring from mothers treated during pregnancy grew faster but were behaviorally similar to control offspring, while offspring from mothers treated during lactation grew slower and were more active in an open-field trial compared to control offspring (Dantzer, van Kesteren, et al., 2020; Westrick et al., 2021). Male offspring from the pregnancy GC treatment also had a greater HPA axis negative feedback response than male offspring in the control group (Westrick et al., 2021).

To follow up on these phenotypic results, in our current study, our goal was to explore the overall impact of elevated maternal GCs on the molecular level. Following the manipulation described above, we tested for differential gene expression in the hypothalamus and hippocampus of the same offspring. While the hypothalamus is a key component of the HPA axis, the hippocampus also has an abundance of glucocorticoid and mineralocorticoid receptors making it considerably sensitive to GCs and allowing it to act as an important inhibitory regulator of the HPA axis (Herman et al., 2003; Maras & Baram, 2012; Mcewen et al., 1968). Studies in animal models and humans suggest that the development of both the hippocampus and hypothalamus are impacted by pre- and postnatal elevations in GCs (Anifantaki et al., 2021; Bale, 2015; Takahashi, 1998). Because we found that the timing of our treatment (pregnancy vs. lactation) impacted the effect of maternal GCs on the physiological and behavioral phenotypes we tested, we were specifically interested in whether the timing of elevated maternal GCs also influenced long-lasting effects on neural gene expression.

## Methods

### Study system

We performed all manipulations and tissue collection between February and August of 2016 and 2017 with a population of North American red squirrels on the traditional land of the Champagne and Aishihik First Nations in the Kluane region of southwest Yukon Territory, Canada (61°N, 138°W). The study population of squirrels lived within a 0.3 km^2^ area. Both sexes of red squirrels are highly territorial and must defend their food cache yearlong to survive overwinter (Dantzer et al., 2012; Hendrix et al., 2020; Siracusa et al., 2017). There are minimal sex differences in the early life history of red squirrels – both sexes must acquire a territory to defend for caching food to survive overwinter, either through dispersal from their natal territory to find their own territory or inheriting their mother’s territory (Berteaux & Boutin, 2000; Boon et al., 2008; Hendrix et al., 2020; Price & Boutin, 1993).

To monitor reproductive status, we live-trapped (Tomahawk Live Trap, Tomahawk, WI, USA) female squirrels and tracked whether fetuses could be felt via abdominal palpation, nipple condition, milk expression, and weight (see Dantzer, McAdam, et al., 2020 for detailed description of trapping methods and long-term study). We ear-tagged female squirrels with unique alphanumeric stamped tags (National Band and Tag Company, Newport, KY, USA) either as a pup (described below) or upon first being trapping as a juvenile or as an adult, if they immigrated from outside the study area. For adults, we used a unique combination of colored wires threaded through each ear tag to identify individuals from a distance. The gestation period in red squirrels is ∼35 days and developing fetuses can be easily felt through abdominal palpation when they are ∼20 days old (Lair, 1985; McAdam et al., 2007). Through frequent trapping around the estimated day of parturition, we monitored mothers’ weight and milk expression, or lack thereof, to identify the date of birth of the litter.

Once parturition was detected, we collared the mother with a VHF radio transmitter (Holohil PD-2C, 4 g, Holohil Systems Limited Carp, Ontario, Canada) and used telemetry to track the mother’s location to the nest with her pups to collect a tissue sample for DNA and collected sex and weight data on individual pups. Pups are born altricial and entirely dependent on their mother’s care (Nice et al., 1956; Svihla, 1930). Approximately 25 days post-parturition, we again used radio telemetry through collaring the mother to find the pups in a nest on the natal territory. At this point, we recorded the weight of each individual again and ear-tagged pups with unique alphanumeric stamped tags (National Band and Tag Company, Newport, KY, USA) in each ear to identify individuals in future recaptures. We used a unique combination of colored washers in each ear tag to be able to identify individual squirrels from a distance after they emerged from the nest around 35-45 days old. At this nest entry, we also collected blood from pups for additional studies (see Dantzer, van Kesteren, et al., 2020).

We conducted all work under the animal care and use approval from University of Michigan (#PRO00005866 and #PRO00007805), as well as Wildlife Research Permits (#0162 and #0205) and Yukon Scientists and Explorers’ Permits (16-19S&E and 17-13S&E) from Yukon Territorial Government.

### Maternal glucocorticoid manipulation

For the glucocorticoid manipulation, we provisioned squirrels with small amounts of peanut butter (∼8 g) and wheat germ (∼2 g) mixed either with or without dissolved hydrocortisone (H4001, Sigma-Aldrich). To mix the GC doses of peanut butter, we first dissolved 800 mg hydrocortisone in 1 mL of 100% ethanol, then mixed the ethanol solution with 5 mL of peanut oil. We let the mixture sit overnight at room temperature to evaporate the ethanol and then combined the peanut oil with 800 g peanut butter and 200 g wheat germ. We weighed out individual doses (∼10 g) and placed each dose in individual containers to be stored at −20°C until needed for provisioning. Each GC treatment peanut butter dose contained 8 mg of hydrocortisone. We selected this dosage to stay within natural physiological levels of GCs based on previous red squirrel studies (Dantzer et al., 2013; van Kesteren et al., 2019). The GC peanut butter treatments rapidly increased plasma cortisol levels after consumption and resulted in a higher cumulative amount of cortisol in the treated females across 24 hours compared to those provided with control treatments (van Kesteren et al., 2019). We made the control doses with the same procedure but without the hydrocortisone in the ethanol and peanut oil mixture. We provisioned squirrels by placing peanut butter treatments in buckets hanging on each squirrel’s territory. We hung the lidded 10.5 L buckets with two holes cut into the side ∼7-10 m off the ground at the center of each territory. Since red squirrels are defensive of their territory, we documented very little to no pilferage of these treatments (van Kesteren et al., 2019).

Once females were confirmed pregnant, we assigned them to one of four treatment groups: pregnancy GC treated, pregnancy control, lactation GC treated, or lactation control. We strategically assigned GC and control treatments across the breeding period to ensure a balanced comparison. All litters were born within 60 days of one another (2016: 60 day range; 2017: 32 day range). Not all litters and not all pups survived to weaning so not all treated litters were included in the current study (for more details on the overall manipulation sample size see Dantzer, van Kesteren, et al., 2020; Westrick et al., 2021). The following information is only for the litters included in the current study. For the pregnancy treatment groups, we aimed to treat mothers for 20 days starting ∼15 days prior to birth (20 days after conception) to until 5 days after birth. Due to variation in detecting the exact date of conception via palpation of fetuses, we actually treated pregnant mothers from 11.8±4.10 days (mean±SD) prior to birth to 4.18±3.11 days after birth for an average total treatment length of 17±3.21 days. We aimed to treat lactating mothers for 10 days starting 5 days post-parturition to allow confirmation of lactation and pup survival. We actually treated lactating mothers from 4.8±1.55 days to 14±1.76 days post-parturition for an average total treatment length of 10.2±0.42 days. In the current study, we sampled offspring from a total of 10 lactation-treated litters (2017: 4 GC litters, 6 control litters) and 25 pregnancy-treated litters (2016: 5 GC, 3 control; 2017: 7 GC, 7 control).

### Tissue dissection

Trapping offspring prior to dispersal from their natal territory as juveniles was essential because, on average, red squirrel survival through their first year of life is low (∼26% survival; Hendrix et al., 2020; McAdam et al., 2007). Juveniles can be more difficult to locate and are at higher risk of dying as they disperse (Hendrix et al., 2020). To catch offspring prior to dispersal, we live-trapped offspring for phenotyping and tissue dissection around weaning at 67.6±3.96 days old (mean±SD; range: 60-81 days old; weaning age is ∼70 days old; Boutin & Larsen, 1993). Due to logistical limitations involved in live trapping, we trapped one to two individuals per litter to maximize the number of litters we could sample at as close to 70 days old as possible. If the litter included both males and females, we aimed to trap one male and one female from the litter if possible (some litters did not include both sexes). Across 2016 and 2017, we collected brain tissue from 16 offspring from the pregnancy-control treatment (2016: 3 male, 2 female; 2017: 3 male, 8 female), and 18 offspring from the pregnancy-GC treatment group (2016: 4 male, 4 female; 2017: 6 male, 4 female). In 2017, we collected brain tissue from 9 offspring from the lactation-control treatment (4 male, 5 female), 5 offspring from the lactation-GC treatment group (2 male, 3 female) -- making a grand total of 48 individual samples across 32 litters.

After trapping, we conducted two behavioral trials (open-field and mirror image stimulation trials) and dexamethasone and adrenocorticotropin releasing hormone (ACTH) challenges on each squirrel for a separate study on behavioral and physiological responses to elevated maternal glucocorticoids (see Westrick et al., 2021 for details). Following the hormone challenges (average of 3.72 hours after dexamethasone injection and 2.58 hours after ACTH injection), we restrained the squirrel in a canvas handling bag and anesthetized with an overdose of isoflurane before rapid decapitation. Importantly, all individuals experienced the behavioral and hormonal experiments. Within 9 minutes after decapitation, we removed the brain, separated the two hemispheres and cerebellum, and froze each section on dry ice. We then kept samples on dry ice until storing at −80° C in the field. We transported samples on dry ice or liquid nitrogen back to Michigan and Illinois and kept samples at ≤ −70° C until microdissection of brain regions in the lab.

We randomly selected one hemisphere from each individual squirrel for microdissection of the hypothalamus and hippocampus. We used a stainless-steel brain matrix and sterile biopsy punches to dissect as much of each region as possible. We sterilized all instruments and tools with RNaseZap® (Ambion, Inc.) and a 100% ethanol rinse prior to dissection. We performed dissections in a cryostat sterilized with 100% ethanol and kept at −22° C to keep the brain frozen during 1mm coronal slicing using the brain matrix and a sterile razor blade. To isolate brain regions, we placed each slice on a sterilized glass slide to visualize both sides of the slice (more rostral and more caudal) to confirm punch accuracy. For instance, if the hippocampus was visible on one side of the slice but not the other, we did not collect a punch from that slice. There is no published brain atlas available for *T. hudsonicus*, therefore to determine the location of each region, we used the thirteen-lined ground squirrel (*Ictidomys tridecemlineatus;* Joseph, Knigge, Kalejs, Hoffman, & Reid, 1966) and rat (*Rattus norvegicus domestica*; Paxinos & Watson, 2006) brain atlases as references. We collected the hypothalamus and hippocampus separately using 1mm sterile biopsy punches resulting in samples of both regions for 48 individuals.

### RNA library construction and sequencing

To extract RNA, we first weighed the tissue samples in 2 mL tubes prefilled with 1.4 mm ceramic beads. We homogenized ≤20 mg samples in 350 µL homogenization buffer and 20-30 mg samples in 600 µL homogenization buffer. We used a QIAGEN TissueLyser® bead mill for 30 seconds at a rate of 20 Hz for two rounds of homogenization. We then used the QIAGEN RNeasy Plus Kit® to extract total RNA following the manufacturer’s instructions. We quantified total RNA concentration on an Invitrogen Qubit® Fluorometer and ran all the samples on agar gels to visualize the 18S and 28S bands.

We sent the total RNA samples for RNAseq library preparation and sequencing at the Roy J. Carver Biotechnology Center at University of Illinois Urbana-Champaign. After assessing RNA quality number on an Agilent Fragment Analyzer, the Biotechnology Center prepared the RNAseq libraries using the Illumina TruSeq Stranded mRNAseq Sample Prep kit, and pooled libraries for sequencing on one S4 lane for 151 cycles on a NovaSeq 6000. Reads were 150 nucleotides in length with an average of 33,688,236 reads per sample (range = 25,075,970 - 49,894,614 reads).

### De novo transcriptome assembly

To assemble *de novo* transcriptomes, we compiled brain-region specific RNA-seq data from four randomly chosen individuals (one from each treatment group: lactation GC, lactation control, pregnancy GC, and pregnancy control). Because we expected baseline differences between brain regions (e.g. Kautt et al., 2024; Nadler et al., 2006), regardless of treatment group, we used Trinity (v2.15.1; Grabherr et al., 2011; Haas et al., 2013) to assemble separate transcriptomes for the hypothalamus and hippocampus. We then used CD-HIT-EST from the program CD-HIT (v4.8.1; Fu et al., 2012; Li & Godzik, 2006) to cluster overlapping contigs and removed any contigs <250bp long. We used Trinotate (v4.0.0; Bryant et al., 2017) to functionally annotate the *de novo* assembled transcriptomes based on homology with known sequence data. Using the same process, we also assembled genome-guided transcriptomes for each brain region using the Eastern grey squirrel (*Sciurus carolensis*) genome (Mead et al., 2020) to compare with our unguided *de novo* assemblies. We ran each assembly through the Benchmarking Universal Single-Copy Orthologue (BUSCO) tool to assess completeness according to conserved orthologs in the Mammalia dataset (v5; Manni, Berkeley, Seppey, Simão, et al., 2021; Manni, Berkeley, Seppey, & Zdobnov, 2021). The guided and unguided transcriptomes both had >80% complete BUSCOs, but the unguided assemblies had slightly higher percentage of single-copy BUSCOs (hippocampus: 49.1% for unguided vs. 44.3% for guided; hypothalamus: 49.3% for unguided vs. 45.3% for guided). We tested the mapping rates of two random samples to the guided and unguided assemblies and found approximately equal percentage of reads mapping. We chose to continue our analyses with the unguided transcriptome assemblies because they had less contigs and a slightly higher percentage of BUSCOs.

The hypothalamus *de novo* transcriptome assembly had 173,835 contigs (83.1% complete BUSCOs; Trinity genes = 132,380; contig N50 = 2.64 kbp; average contig length = 1.24 kbp) and the hippocampus *de novo* transcriptome assembly had 180,728 contigs (81.9% complete BUSCOs; Trinity genes = 137,560; contig N50 = 2.64 kbp; average contig length = 1.23 kbp). We performed all subsequent analyses separately for the hippocampus and hypothalamus using the respective transcriptome assembly. Our final, annotated *T. hudsonicus* transcriptome assembly is available on [*to be updated upon acceptance*]. Raw reads are available from the NCBI SRA [*to be updated upon acceptance*].

### Differential expression analysis

For each sample, we estimated transcript abundance using Salmon (v1.10.1; Patro et al., 2017). We used Salmon to produce a matrix of estimated transcript counts for downstream analysis in R (version 4.3.1; R Core Team, 2023). We first filtered for genes with ≥ 10 reads across all samples. Using DESeq2 (M. I. Love et al., 2014), we next normalized counts at the ‘gene’ level and quantified transcript enrichment or depletion in a pair-wise fashion as a log-fold difference between GC-treated and control groups for both pregnancy and lactation. Given known differences in neural gene expression between male and female individuals (Gegenhuber & Tollkuhn, 2020; McCarthy et al., 2009), we included sex as a main effect in the model to additionally look for transcript enrichment or depletion between female and male offspring. While previous studies on maternal stress have found sex differences in the effect on offspring (e.g. Metzger & Schulte, 2016), we were underpowered to test for an interaction between GC treatment and sex due to small sample sizes, particularly in the lactation treatment group. After p-value adjustment for multiple comparisons with the Benjamini-Hochberg correction (Benjamini & Hochberg, 1995), we used a false discovery rate (FDR) cutoff of 0.1 to identify significantly differentially expressed genes. To make the heatmaps for visualizing the relationships between samples, we used a variance stabilizing transformation in DESeq2 and used the R package pheatmap (Kolde, 2019). To visualize overall expression variance, we performed a principal components analysis (PCA) of all transcript expression values after the variance stabilizing transformation using the plotPCA function in DESeq2. Given their importance in regulating the HPA axis and their proposed role in maternal stress effects on offspring (Hellstrom et al., 2012; Liu et al., 1997; Yehuda et al., 2016; Zimmer et al., 2021), we also specifically looked for differential expression of the transcripts for the FK506-binding protein, glucocorticoid receptor (GR), and mineralocorticoid receptor (MR).

Using the R package RRHO2 (Huo, 2016), we looked for patterns of concordance in gene regulation to assess whether the pregnancy and lactation treatments had similar patterns in gene expression. The threshold-free rank-rank hypergeometric overlap (RRHO) algorithm looks for statistical significance in the overlap of two ranked gene lists. We compared gene lists ranked by degree of differential gene expression for the pregnancy and lactation treatment in each brain region separately. RRHO analysis helps us understand how similar overall gene expression changes are between the pregnancy and lactation treatments independent of the statistical significance of individual genes. We performed the RRHO analysis across all genes with non-zero counts in both the pregnancy and lactation treatments.

### Functional enrichment analysis

To investigate the functional impact of differential gene expression, we ran gene ontology (GO) enrichment analysis for each brain region. GO analysis asks which biological processes are over- or under-represented given an annotated list of differentially expressed genes. With the R package topGO (Alexa & Rahnenfuhrer, 2023), we generated the GO graph structure of biological processes with the “weight01” algorithm and used Fisher’s exact test to identify enriched terms.

## Results

### Hippocampus

There were 129,633 total genes detected with greater than 10 reads across all samples in the hippocampus pregnancy dataset. Out of these 129,633 genes, we found 9 genes were upregulated (0.0069%) and 76 genes were downregulated (0.059%) in the pregnancy GC treated offspring compared to the control offspring (Figure 1; for full list of genes see Supplemental Materials Table S1). Out of 129,633 total genes across pregnancy treatment groups, we found 40 upregulated genes (0.031%) and 176 downregulated genes (0.14%) in the hippocampus of female offspring compared to male offspring (Figure 2; Supplemental Materials Table S2). In a PCA of data across treatment groups, the first principal component explained 11% of the variance in and the second component explained 8% of the variance in overall gene expression (Figure 1 and Figure 2).

**Figure 1.**
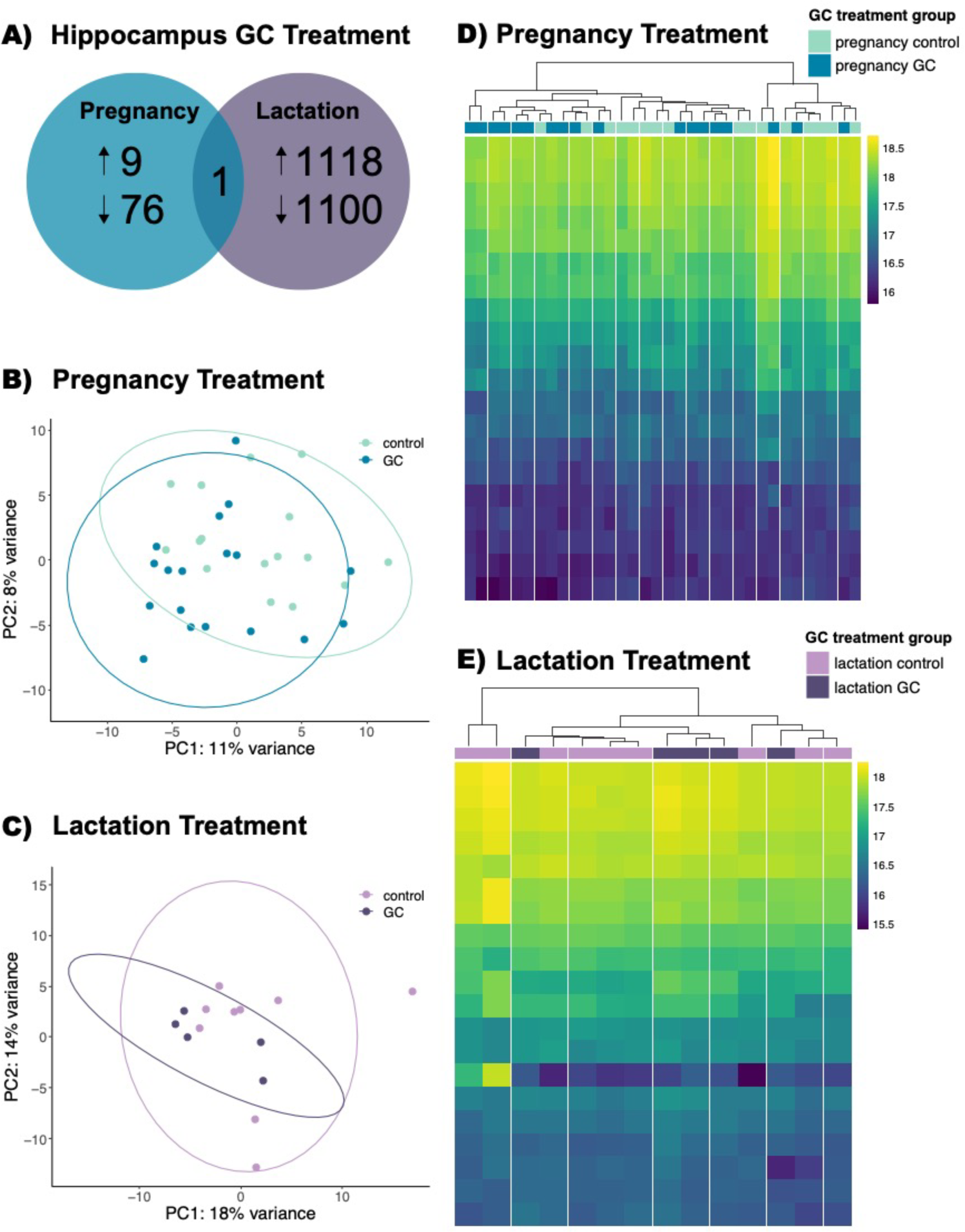
Differential gene expression in the hippocampus. (A) We found 9 up-regulated and 76 down-regulated genes in the pregnancy GC treatment group compared to the pregnancy control group. We found 1118 up-regulated and 1100 down-regulated genes in the lactation GC group compared to the lactation control group. Only one gene was in common between the gene sets as illustrated by the overlap of the Venn diagram. (B) In a principal component analysis across all gene expression data from the pregnancy treated individuals, the first component explained 11% of the variance and the second component explained 8% of the variance in gene expression. The lighter blue color represents the pregnancy control group, and the darker blue color represents the pregnancy GC group. (C) In a principal component analysis across all gene expression data from the lactation treated individuals, the first component explained 18% of the variance and the second component explained 14% of the variance in gene expression. The lighter purple color represents the lactation control group, and the darker purple color represents the pregnancy GC group. These heatmaps show normalized transcript counts (rows) clustered based on expression similarity of individual samples (columns) across groups for the top 20 most variable genes from the (D) pregnancy treatment time period and (E) lactation treatment time period. Columns are labeled at the top with the color of their respective treatment group.

**Figure 2.**
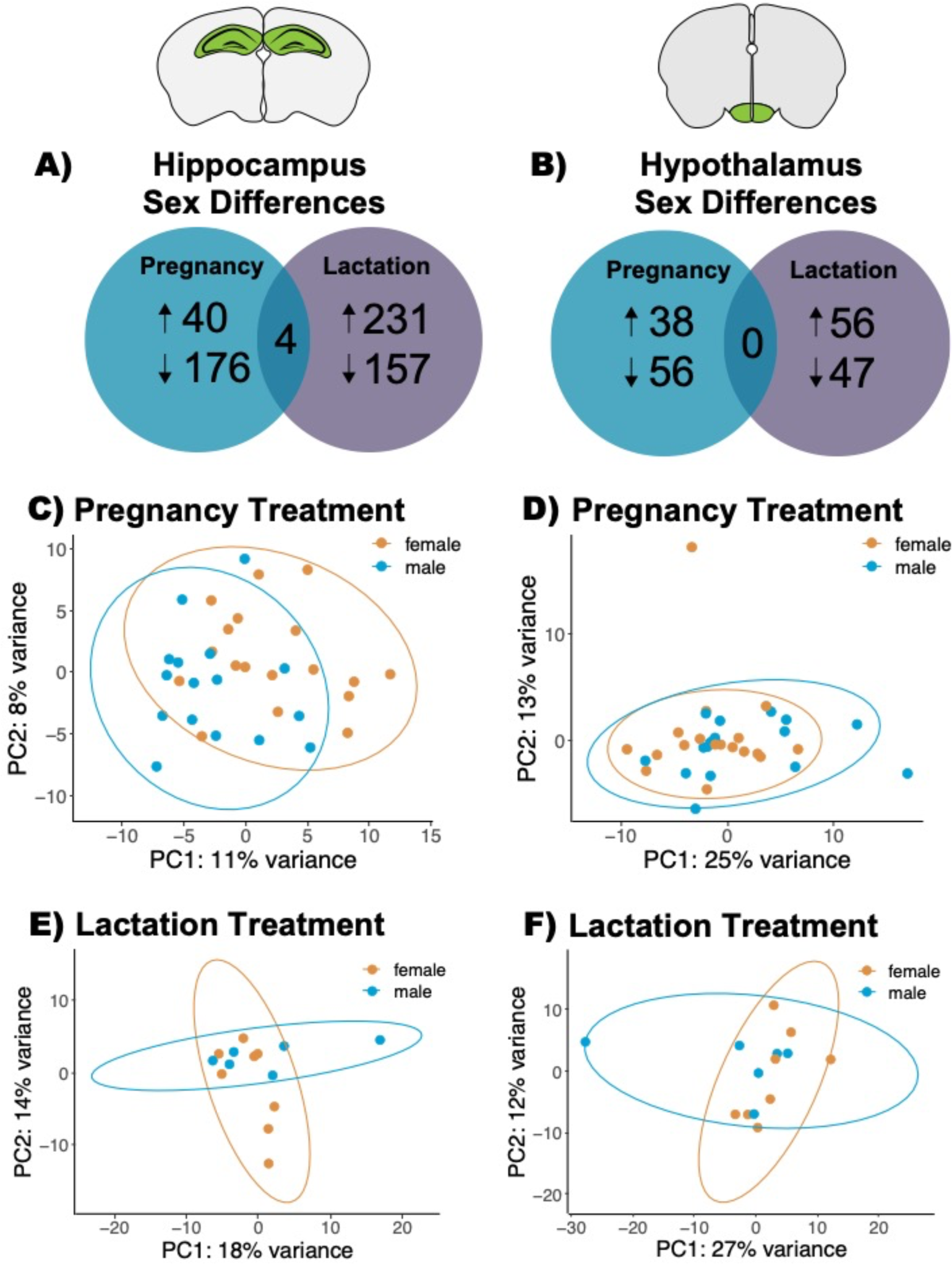
Differential gene expression between female and male offspring. (A) When controlling for treatment group (GC vs control), we found 40 up-regulated and 176 down-regulated genes in the hippocampus of female offspring from the pregnancy treatment group compared to male offspring. In the hippocampus of female offspring from the lactation treatment group, we found 231 up-regulated and 157 down-regulated genes compared to male offspring. There were four genes in common between differentially expressed gene sets. (B) In the hypothalamus of female offspring from the pregnancy treatment group, we found 38 up-regulated genes and 56 down-regulated genes compared to male offspring. We found 56 up-regulated genes and 47 down-regulated genes in the hypothalamus of female offspring from the lactation treatment group compared to male offspring. (C) In a principal component analysis of the hippocampus data from the pregnancy treatment, the first component explained 11% of the variance and the second component explained 8% of the variance in gene expression profiles. (E) The first component explained 18% of the variance and the second component explained 14% of the variance in gene expression profiles in the hippocampus of offspring from the lactation treatment group. In principal component analyses of the hypothalamus data, (D) the first component explained 25% of the variance and the second component explained 13% of variance in gene expression profiles from offspring in the pregnancy treatment group. (F) In the lactation treatment group, the first component explained 27% of variance and the second component explained 12% of variance in gene expression profiles. In both Venn diagrams (A,B), the blue color indicates data from the pregnancy treatment and the purple color indicates data from the lactation treatment. Across all principal component scatterplots (C-F), the orange color indicates female offspring, and the blue color indicates male offspring.

There were 22,354 total genes with greater than 10 reads across all samples in the lactation treatment dataset. Out of these 22,354 total genes, we found 1118 genes were upregulated (5%) and 1100 genes were downregulated (4.9%) in the lactation GC treatment offspring compared to the control offspring (Figure 1; for full list of genes see Supplemental Materials Table S3). Out of 111,368 genes with nonzero total read count, we found 231 upregulated genes (0.21%) and 157 downregulated genes (0.14%) in the hippocampus of female offspring compared to male offspring (Figure 2; Supplemental Materials Table S4). In a PCA of data across treatment groups, the first principal component explained 18% of the variance and the second component explained 14% of the variance in overall gene expression (Figure 1 and Figure 2).

Between the pregnancy and lactation treatment time periods, we found one gene was differentially expressed in both pregnancy and lactation GC treatments compared to control (Figure 1). Four genes were differentially expressed in both pregnancy and lactation-treated females compared to males (Figure 2). Since we saw few statistically significant genes overlapped between the two treatment time periods, we used RRHO analysis to understand general patterns across all genes regardless of statistical significance. Overall, our RRHO analysis showed strong concordance between the effect of pregnancy and lactation treatments in the regulation of genes (Figure 4), meaning both GC treatments had similar impacts in the direction of gene regulation. The strongest concordance between treatment groups (pregnancy and lactation) was in the downregulation of genes in the GC-treated offspring compared to control (Figure 4).

Because there were few differentially expressed genes from the pregnancy treatment (85 genes), we ran GO term analysis on the differentially expressed genes in the lactation treatment dataset only (2218 genes). For the lactation treatment, GO term enrichment analysis indicated over-representation of genes associated with processes including cytoplasmic translation, energy coupled protein transport, regulation of glutamate receptor signaling, and positive regulation of axonogensis (see Supplemental Materials Table S5 for full list). Notably, ‘cellular response to glucocorticoids’ was not among the top 100 over-represented GO terms, though ‘glucocorticoid metabolic process’ was the 36^th^ over-represented GO term.

Given our *a priori* selection of genes for proteins essential to the HPA axis (GR, MR, and FKBP5), we also looked at GR, MR, and FKBP5 genes without FDR correction. In the lactation treatment group, we found pups whose mothers were treated with GCs exhibited decreased gene expression of the MR (log2foldchange = −-0.22, p = 0.02) and female offspring expressed higher levels of GR than male offspring (log2foldchange = 0.18, p = 0.01). However, we did not find significant differences in expression of GR, MR, and FKBP5 between the GC and control groups for either treatment (pregnancy or lactation) based on the adjusted p-value (p>0.1).

### Hypothalamus

After filtering, out of 127,247 total genes detected in the hypothalamus pregnancy dataset, we found four genes were upregulated (0.0031%) and one gene was downregulated (0.000079%) in the pregnancy GC treated offspring compared to the control offspring (Figure 3; for full list of genes see Supplemental Materials Table S6). Across pregnancy treatment groups, out of the 54,301 genes included in the sex comparison, 38 genes were upregulated (0.07%) and 56 genes were downregulated (0.1%) in the hypothalamus of female offspring compared to male offspring (Figure 2; for full list of genes see Supplemental Materials Table S7). In a PCA of data across treatment groups, the first principal component explained 25% of the variance and the second component explained 13% of the variance in overall gene expression (Figure 2 and Figure 3).

**Figure 3.**
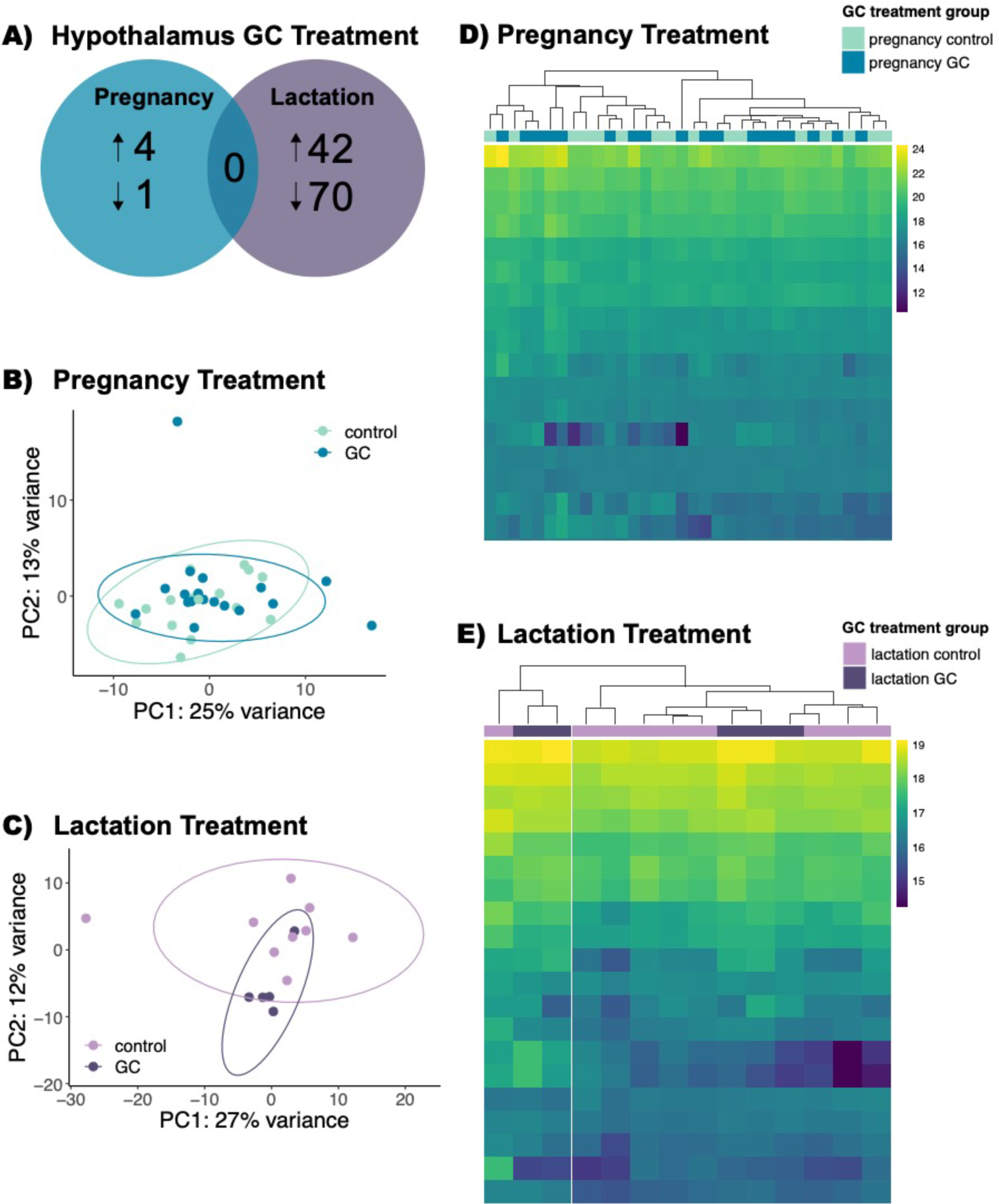
Differential gene expression in the hypothalamus. (A) We found 4 up-regulated and 1 down-regulated genes in the pregnancy GC treatment group compared to the pregnancy control group. We found 42 up-regulated and 70 down-regulated genes in the lactation GC group compared to the lactation control group. No genes were in common between the gene sets as illustrated by the overlap of the Venn diagram. (B) In a principal component analysis across all gene expression data from the pregnancy treated individuals, the first component explained 25% of the variance and the second component explained 13% of the variance in gene expression. The lighter blue color represents the pregnancy control group, and the darker blue color represents the pregnancy GC group. (C) In a principal component analysis across all gene expression data from the lactation treated individuals, the first component explained 27% of the variance and the second component explained 12% of the variance in gene expression. The lighter purple color represents the lactation control group, and the darker purple color represents the pregnancy GC group. These heatmaps show normalized transcript counts (rows) clustered based on expression similarity of individual samples (columns) across groups for the top 20 most variable genes from the (D) pregnancy treatment time period and (E) lactation treatment time period. Columns are labeled at the top with the color of their respective treatment group.

Out of 94,841 total genes in the hypothalamus lactation dataset, we found 42 genes were upregulated (0.044%) and 70 genes were downregulated (0.074%) in the GC treated offspring compared to the control offspring (Figure 3; for full list of genes see Supplemental Materials Table S8). Across lactation treatment groups, out of the 84,910 genes included in the sex comparison, 56 genes were upregulated (0.066%) and 47 genes were downregulated (0.055%) in the hypothalamus of female offspring compared to male offspring (Figure 2; for full list of genes see Supplemental Materials Table S9). In a PCA of data across treatment groups, the first principal component explained 27% of the variance and the second component explained 12% of the variance in overall gene expression (Figure 2 and Figure 3).

In the hypothalamus, we did not find any overlap in differentially expressed genes between treatment time periods for the GC treatment or sex (Figure 2 and Figure 3). Despite this lack of overlap, our RRHO analysis showed concordance between pregnancy and lactation treatment groups in the differential expression of genes (Figure 4). The strongest concordance between treatment groups in gene regulation was in the upregulation of genes (Figure 4).

**Figure 4.**
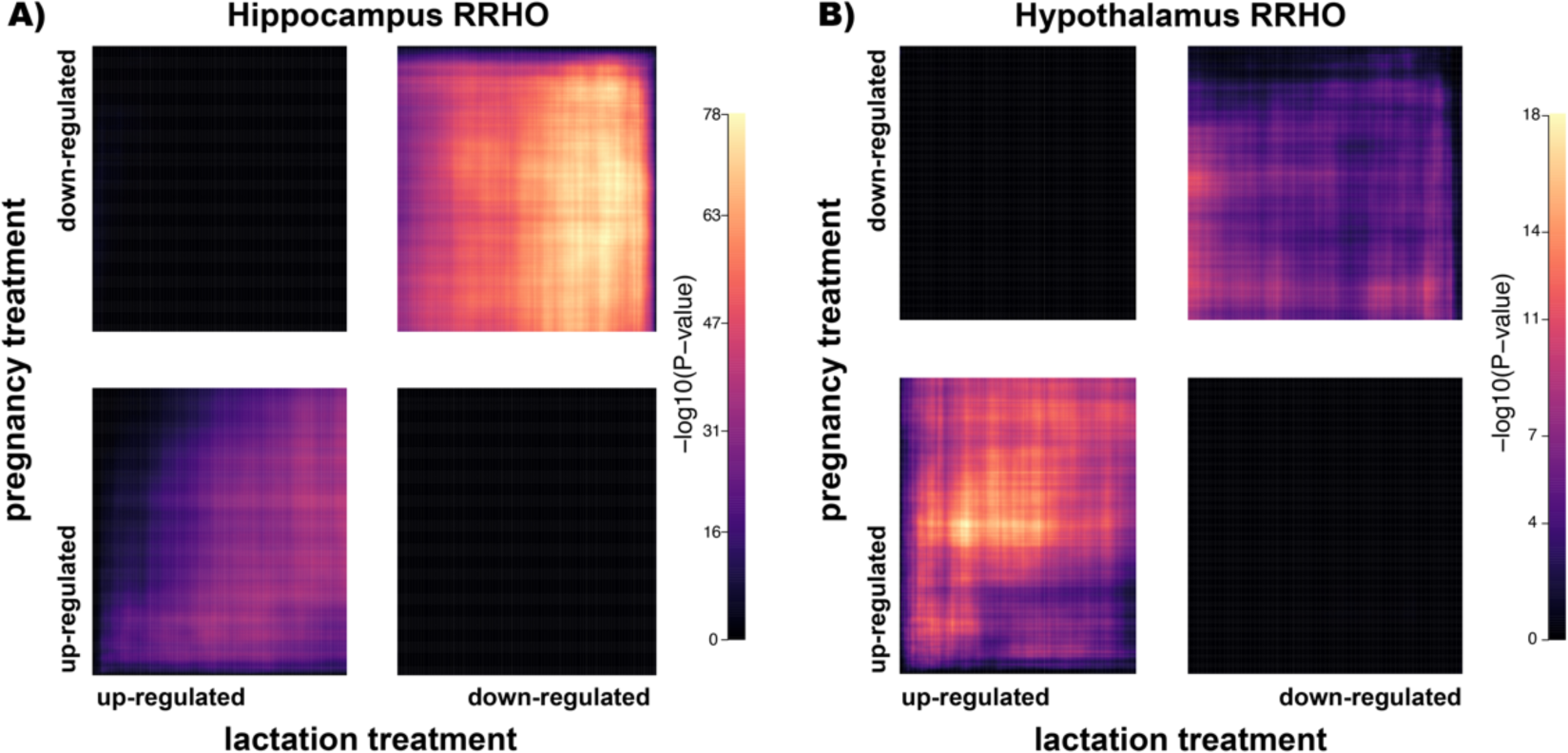
Rank-rank hypergeometric overlap analysis. Each panel is broken into four regions: up-regulated in both treatment groups (bottom left), down-regulated in both treatment groups (top right), up-regulated in the pregnancy treatment but down-regulated in the lactation treatment (top left) and down-regulated in the lactation treatment but up-regulated in the pregnancy treatment group (bottom right). The color corresponds to the value of the −log10(p-value) from the differential expression analysis. Warmer colors (hot spots) indicate the strength and directionality of concordance in gene expression between treatment groups. Genes from the hippocampus analysis are in panel (A) and genes from the hypothalamus analysis are in panel (B).

Given the small number of differentially expressed genes in the pregnancy treatment (5 genes), we again only ran GO term analysis on the lactation treatment group. Across the genes that were differentially expressed between lactation treatment groups (112 genes), the top 10 GO terms enriched included T cell receptor signaling pathway and positive regulation of natural killer cell differentiation (for full list of GO terms see Supplemental Material Table S10). Again, cellular response to glucocorticoids and regulation of glucocorticoid mediated signaling were notably not among the top 100 over-represented GO terms. Additionally, we did not find significant differences in expression of the genes for FKBP5, GR, and MR between the GC and control groups or between females and males of either treatment period.

## Discussion

Maternal stress can have important life-long effects on offspring physiology and behavior (Bale, 2005; Barbazanges et al., 1996; Glover et al., 2010), which are thought to be mediated through epigenetic changes altering gene expression (Reshetnikov et al., 2018; Weaver et al., 2002). While the impact of maternal stress on gene expression in offspring has been well studied in laboratory species (e.g., Peña et al., 2019), how these effects vary with the timing of maternal stressors and whether the effects are detectable is less clear in natural populations, where they have important consequences for evolution. Our goal in this study was to investigate whether maternal stress has long-lasting influences on offspring at a molecular level in a free-ranging mammal. Specifically, we were interested in the impact of maternal GCs experienced *in utero* or through early lactation on neural gene expression in offspring at independence. After feeding pregnant or lactating mothers GCs, we trapped offspring at weaning to measuring RNA expression in the hypothalamus and hippocampus, given the importance of both brain regions in the physiological stress response (Charmandari et al., 2005; Tsigos & Chrousos, 2002) and behavior (Behrendt, 2011; Goodson, 2005).

Overall, we found differences in gene regulation between the different treatment time periods tested across both brain regions. We found more genes were both up- and down-regulated between GC-treated and control offspring in the lactation treatment compared to the pregnancy treatment in both the hippocampus and hypothalamus. Though the qualitative pattern was the same across both brain regions, we found more genes were significantly differentially expressed in the hippocampus (2218 genes, 9.9% up- or down-regulated) than the hypothalamus (114 genes, 0.12% up- or down-regulated) in the lactation treatment group. Across both brain regions, we found strikingly few differentially expressed genes overlapping between the two treatment time periods (pregnancy and lactation) suggesting variation in gene targets based on timing of maternal GC exposure. This result is consistent with our behavioral and physiological results from the same offspring which indicate more behavioral differences in offspring of the lactation treatment than the pregnancy treatment (Table 1; Westrick et al., 2021). Taken together, these results provide further evidence for how the timing in development can impact the consequences of the same early life experiences.

**Table 1.**
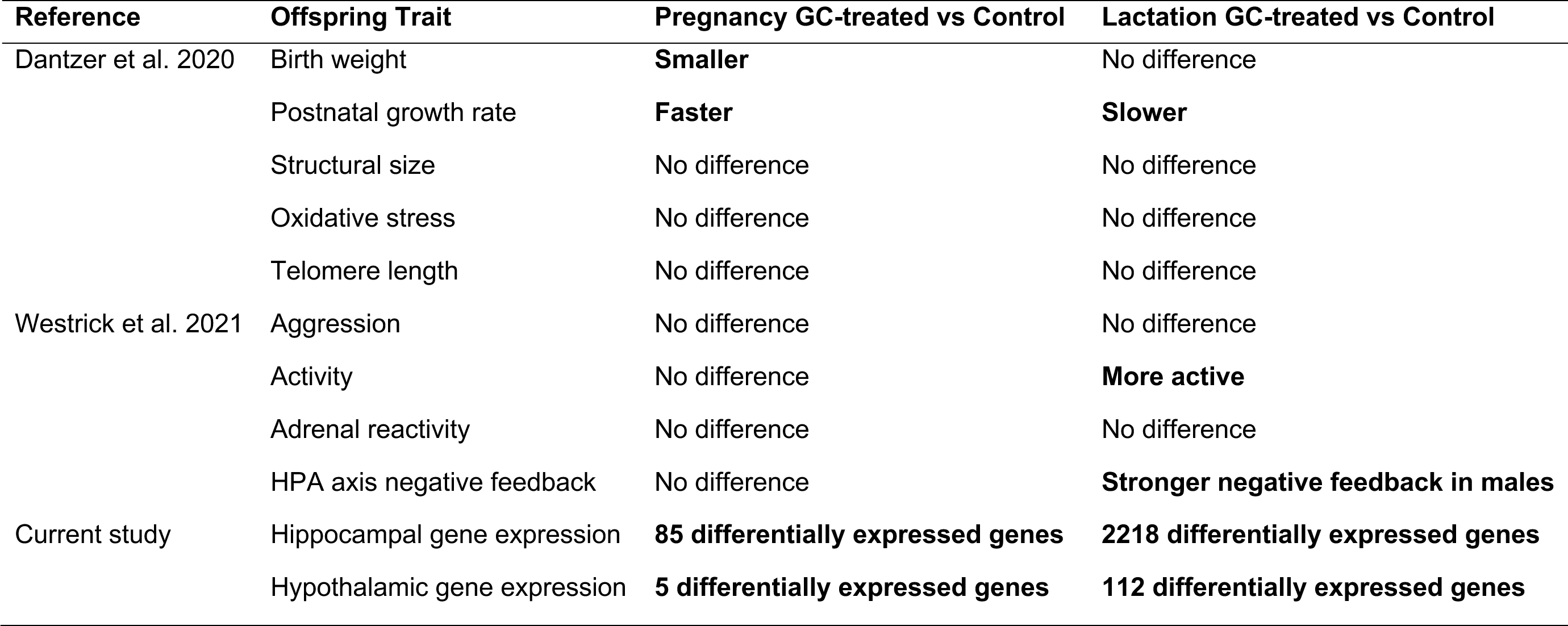
Summary table of results from maternal glucocorticoid manipulation.

There are a few non-mutually exclusive hypotheses for why the postnatal (lactation) treatment showed a stronger impact on neural gene expression than the prenatal (pregnancy) treatment. We hypothesize that offspring may be more buffered from maternal GCs during pregnancy and/or offspring may be more sensitive to maternal GCs during lactation. To begin with, the placenta plays an important role of buffering offspring *in utero* from maternal GCs (Seckl & Holmes, 2007; Shearer et al., 2019; Sze & Brunton, 2024). Given this buffering effect, it is possible that increasing maternal GCs during pregnancy did not significantly impact fetal circulating GC levels. However, past work has shown injecting pregnant rats with corticosterone results in higher levels of circulating corticosterone in pups immediately after birth (Brummelte et al., 2010). Additionally, we do see a significant increase in early growth rate of squirrel pups from the pregnancy GC treatment (Table 1; Dantzer et al., 2013; Dantzer, van Kesteren, et al., 2020), which indicates the offspring are not entirely buffered from impacts of an increase in maternal GCs during pregnancy. Though this change in postnatal growth rate may not necessarily reflect increased transmission of maternal GCs during pregnancy to offspring, but offspring growth may potentially be impacted indirectly through a change in maternal behavior (e.g. Nephew & Bridges, 2011; Nguyen et al., 2008).

Furthermore, in many mammals, neural development continues during lactation (Watson et al., 2006) and therefore, the lactation treatment may have been during a critical time window (sensitive period), impacting the development of the hippocampal dentate gyrus in particular (Watson et al., 2006). This may also explain why the offspring from the lactation GC treatment group show more behavioral differences from control than pregnancy GC treatment (Westrick et al., 2021), and why the hippocampus showed more differences between treatment groups in gene expression. Alternatively, offspring could be responding indirectly to GC treatment, if downstream treatment effects in mothers (e.g. changes in maternal behavior) depend on treatment timing. In other words, timing dependent differences in offspring could be direct or indirect effects of GCs, and these alternatives are non-mutually exclusive. In either case, our use of GC treatment pinpoints these effects as compared to those of complex, environmental stressor that may also act via alternative mechanisms. Future studies investigating windows of sensitivity to parental cues, such as maternal GCs, will improve our understanding of the mechanisms underlying developmental and transgenerational plasticity (Bell & Hellmann, 2019). In the wild, these sensitive windows may play an important role in shaping offspring phenotypes if they experience considerable variation in stressors across development (e.g. seasonal variation in food availability).

Despite differences between treatment groups in the number of significantly differentially expressed genes, p-value independent RRHO analysis revealed the qualitative overall pattern of gene regulation is consistent across treatment groups. In other words, the pregnancy GC treatment changed the transcriptomic profile in the same direction as the lactation GC treatment in both the hypothalamus and hippocampus, but the lactation treatment had a stronger effect on individual genes. Interestingly, the directionality of the concordance is reversed between the hypothalamus and hippocampus with a stronger pattern of concordant upregulation of genes in the hypothalamus and more concordant downregulation of genes in the hippocampus. These results suggest there is consistency in how elevated maternal GCs impact neural gene expression independent of timing; however, treating mothers during lactation produced a stronger response perhaps due to a greater degree of buffering in offspring from the pregnancy treatment group. These differences between the pre- and postnatal maternal GC treatments suggest variation in when mothers experience stressors can shape the long-term consequences for offspring.

We did not find a large effect of maternal GCs on neural gene expression in terms of the proportion of genes differentially expressed. This result mirrors our previous findings of minimal effects of exogenous maternal GCs on offspring behavior and HPA axis stress physiology at weaning (Table 1; Westrick et al., 2021). While exogenous maternal GCs do impact postnatal growth rate (Table 1; Dantzer, van Kesteren, et al., 2020), because we collected brain samples at weaning (∼55 days post-lactation treatment and ∼65 days post-pregnancy treatment), it is possible that any impacts of the maternal treatments were compensated for throughout further development following the treatments. Our results indicate there are few long-lasting impacts of maternal GCs on the neurogenomics of juvenile red squirrels suggesting compensatory mechanisms to buffer offspring as they develop. While offspring can receive important information about the environment through maternal cues, they may also buffer their responses to avoid the potential costs of developmental plasticity. Juveniles may be prioritizing immediate cues received, rather than maternal cues, in shaping the development of adult phenotypes (McNamara et al., 2016). Future studies should investigate whether juveniles compensate for a mismatch between early life “stress” levels and experiences upon emerging from the nest or if the impact of maternal GCs on neural gene expression patterns is consistent across development.

Even though there are minimal sex differences the early life behavior of juvenile red squirrels, we did find differences in neural gene expression between male and female offspring in both brain regions. The region with the greatest number of differentially expressed genes based on sex was the hippocampus. Specifically, we found offspring from the lactation treatment (both control and GC-treated) had more differentially expressed genes between female and male offspring (0.35% up- or down-regulated genes) in the hippocampus when compared to the pregnancy treated offspring (0.17% up- or down-regulated genes). In the hypothalamus, we found a similar percentage of genes were differentially expressed in female versus male offspring across both treatment time points (pregnancy: 0.17%, lactation: 0.12% up- or down-regulated genes). In other words, we find the largest effect of sex (female vs male) in the hippocampus of offspring from mothers treated during lactation. Across both brain regions, the lack of substantial overlap between gene sets of differentially expressed genes due to sex suggests we are not detecting general sex differences but rather differential tuning of gene expression in both sexes based on their early life experience which could indicate sex-specific effects of maternal GCs. Given both sexes experience the same challenges to survival in their first year (e.g. predation risk, territory acquisition), we might predict few sex-specific differences in the impact of maternal GCs on phenotypes expressed during this year. However, daughters may be able to somewhat mitigate these challenges through being more likely than sons to inherit their territory from their mothers (Fisher et al., 2017), thus eliminating the risks associated with dispersal. Alternatively, given overall differences in neural development of males and females (Lenz et al., 2012; McCarthy et al., 2009), maternal GCs may play a different role in neural development between the sexes. With our limited sample size, we did not explore the interaction between sex and GC treatment, but this is an exciting future avenue of research.

The natural environment the squirrels lived in brought valuable complexity to our study. While all squirrels in this study lived relatively close to one another and experienced roughly the same environmental variation in food supply and predation risk, mothers and offspring likely experienced variable environmental stressors during the experiment. Additionally, individuals vary in how they perceive or detect external stressors and vary in their internal condition. This variation may have diluted or buffered any potential outcomes of our experiment. For example, prenatal stress can alter the consequences of postnatal stress (e.g. Lehmann et al., 2000) and postnatal stress can improve deficits due to prenatal stress (e.g. Crombie et al., 2021). We also note that there can be significant among-year variation in external stressors in this system as predation risk, food availability, and conspecific competition all cyclically fluctuate (Dantzer, McAdam, et al., 2020; Petrullo et al., 2022). Our lactation treatment samples are from one year (2017) and our pregnancy treatment samples are across two years (2016 and 2017). Neither of these years was a white spruce mast year or the year following a mast, which are considered highly stressful years compared to non-mast years (Dantzer et al., 2013; Petrullo et al., 2023). Thus, these two years were relatively similar in environmental conditions. Yet through all of this variation, we still detected an effect of exogenous maternal GCs on neural gene expression suggesting that maternal effects can have robust persistent impacts on neurogenomics in a wild mammal. There are many challenges to studying the mechanisms of maternal effects in the wild, but these challenges are worth tackling to gain valuable insight into how natural populations cope with stressors and how maternal effects shape evolutionary processes.

In conclusion, timing during early life is an important variable in the impact of maternal GCs at a molecular level. Whether due to buffering or differences in sensitive periods of development, the same maternal cue of increased GCs can have different long-term consequences on offspring depending on whether offspring received the cue while *in utero* or through milk. Additionally, we found the same cue influences brain regions in a unique way, underscoring the need to examine brain regions separately rather than assessing brain gene expression across the brain as a whole (Fischer et al., 2021). Given the fitness implications of maternal effects, more studies are needed exploring the mechanisms underlying maternal effects in wild populations. While studies in laboratory populations have been vital in developing predictions about the mechanisms of maternal effects, investigating these questions in wild populations will allow us to assess their actual impact in a natural context. By studying free-ranging mammals, we can return to the roots of neuroethology and develop the tools to bridge across disciplines in our understanding the neuroendocrine mechanisms shaping evolutionary trajectories (Laurent, 2020; Zilkha et al., 2016).

## Acknowledgements

We thank Agnes MacDonald, her family, and the Champagne and Aishihik First Nations for allowing us to conduct our work within their traditional lands. We thank Monica Cooper, Zach Fogel, Claire Hoffmann, Noah Israel, Sean Konkolics, Laura Porter, Matt Sehrsweeny, Sam Sonnega, Jess Steketee, and Dylan Yaffy for their assistance in the field experiment and data collection. This work was supported by the American Society of Mammalogists (SEW), the University of Michigan (SEW and BD), the University of Illinois Urbana-Champaign (EKF), National Science Foundation (IOS-1749627 to BD, PRFB #2010714 to SEW), and the Natural Sciences and Engineering Research Council of Canada (SB, AGM, and JEL). This is publication #[*to be updated upon acceptance*] of the Kluane Red Squirrel Project.

## Data accessibility

Our *T. hudsonicus* transcriptome assembly is available on FigShare https://figshare.com/s/a35ac5f5bb5c1e9abfe1 [*note: URL to be updated upon acceptance, will upload on NCBI as well*]. Raw reads are available from the NCBI SRA (*to be updated upon acceptance*). Relevant metadata and code for the analysis are available on FigShare https://figshare.com/s/a35ac5f5bb5c1e9abfe1 [*note: URL to be updated upon acceptance*].

## Author contributions

SEW and BD designed the study. SEW, FvK, and BD conducted the field work. SEW conducted all laboratory work and performed all analyses under the guidance of EKF. SEW wrote the original manuscript draft. All authors contributed to the review and editing of the manuscript. SEW, SB, AGM, JEL, and BD contributed funding for this research.

